# Exaggerated estrous swellings and male mate choice in primates: testing the reliable indicator hypothesis in the Amboseli baboons

**DOI:** 10.1101/007278

**Authors:** Courtney L. Fitzpatrick, Jeanne Altmann, Susan C. Alberts

## INTRODUCTION

The evolution of male mate choice in sex-role reversed species can easily be understood according to the general principles of sexual selection. That is, when females are mate-limited they are expected to compete for access to males and those males, in turn, are expected to be choosy. This phenomenon is well described in some sex-role reversed species of pipefish (Berglund et al. 1989; Vincent et al. 1992) and shorebirds (Szekely & Reynolds 1995). In contrast, when the sex roles are conventional, choosy males will be at a competitive disadvantage and, therefore, male mate choice should be selected against. Of course, exceptions to this prediction have been demonstrated, most notably in those cases where female fecundity varies and choosy males therefore benefit by selecting females with higher fecundity (Servedio & Lande 2006). Indeed, empirical work across a broad range of taxa indicates that male choosiness in these scenarios is common. For example, in those species where female body size predicts fecundity, males often demonstrate a preference for larger females (lizards; Olsson 1993, fish; Sargent et al. 1986, insects; Bonduriansky 2001).

However, male mate choice is increasingly observed even when direct fecundity benefits for choosy males are not apparent. In some of these cases, males might receive indirect benefits from females, analogous to the “good genes” models of mate choice that have been developed for female choice (see Chapter 3 Andersson 1994) For example, experimental work in barn owls suggests both that male barn owls prefer females with spottier plumage and that the offspring of those females mount a more effective immune response (Roulin et al. 2000, 2001). That is, males may bias their mating investment toward females whose offspring have superior genetic immunity. In other cases, males may evolve preferences when male investment in mating is so high or costly that mating necessarily limits future mating opportunities, thus imposing opportunity costs. For example, males should choose mates carefully when their ability to mate with many females is constrained by intense male-male competition or sperm competition (Schwagmeyer and Parker 1990; Owens and Thompson 1994; reviewed by Edward and Chapman 2011).

The exaggerated estrous swelling displayed by many cercopithecine primates is commonly cited as a trait upon which males base mate choice, and studies in several species have suggested that males prefer females with larger swellings (Domb & Pagel 2001; Higham et al. 2009; Huchard et al. 2009). However, the fitness benefits that males might receive as a result of this choosiness based on swelling size are unknown. Therefore, knowledge of the evolutionary forces that have shaped both the behavior and the trait remain obscured, and the question of what information males may receive from these signals remains open (Deschner et al. 2004; Higham et al. 2009; Huchard et al. 2009; Fitzpatrick et al. 2014).

Sexual swellings appear during the follicular phase of the female sexual cycle, are thought to have evolved multiple times in the primate lineage (Dixson 1983), and are most commonly associated with multi-male multi-female social systems (Nunn 1999). It has been well established in many species that male mating behavior increases in response to these changes in swelling size within a given cycle (baboons, Hausfater 1975; Packer 1979; Bielert & Anderson 1985; Higham et al. 2008b; Huchard et al. 2009; Nitsch et al. 2011; chimpanzees, Tutin 1979; Emery & Whitten 2003; Deschner et al. 2004; Breaux et al. 2012 macaques, van Noordwijk 1985; Brauch et al. 2007). In fact, as the understanding of this trait has become more refined over the past decade, researchers have identified three distinct types of variation in swelling size (Zinner et al. 2002).

1. Swelling size varies within a cycle, generally achieving maximal size for that cycle around the time of ovulation. This type of variation has been demonstrated repeatedly to be a probabilistic indicator of ovulation in at least some species (Hendrickx and Kraemer 1969; Wildt et al. 1977; Dahl et al. 1991; Emery and Whitten 2003; see Nunn 1999; Alberts and Fitzpatrick 2012 for thorough review) and it has been shown in baboons that females are more attractive to males during the period of highest fertility (Noe & Sluijter 1990). Furthermore, it has been demonstrated in chimpanzees that males both copulate with and compete for females more actively during the peri-ovulatory period (Deschner et al. 2004).
2. Swellings vary in size within an individual across cycles. That is, females of most primate species cycle repeatedly before conceiving and it has been demonstrated in several species that maximal swelling size increases from cycle to cycle within one individual (Emery and Whitten 2003; Deschner et al. 2004; Higham et al. 2008b; Fitzpatrick et al. 2014). If the probability of conception increases with subsequent cycles, this type of variation may signal differences in the probability of conception across cycles. Indeed, although variation in conception probability across cycles within an individual has not—to our knowledge—been described empirically, data from our study population suggest that the cycles immediately following post partum amenorrhea are less likely to result in conception than later ones (data presented in Fitzpatrick et al. 2014.) Furthermore, it has been shown that alpha males successfully target conceptive cycles, suggesting that, within a given female, cycles vary in the probability of conception (Weingrill et al. 2003; Alberts et al. 2006).
3. Finally, swelling sizes may differ between individuals. Because this type of variation can only be revealed by controlling for the within-cycle and across-cycle variation exhibited by each female, it is more challenging to demonstrate empirically. Three studies have been able to do so (Deschner et al. 2004; Huchard et al. 2009; Fitzpatrick et al. 2014).

Thus, it is known that males respond to the first type of variation (within-cycle variation in swelling size) and that the second type of variation has the potential to signal important information to males (because swelling size increases as cycle number progresses). However, it remains unclear whether the third type of variation—variation between females in sexual swelling size (beyond differences accounted for by within-cycle and across-cycle variation)—has additional information content for male primates.

Despite this gap, a main hypothesis that attempts to explain both the evolution of exaggerated swellings and the male response to them hinges on the assumption that it is precisely this variation between individuals that is salient for males. This hypothesis has become known as “the reliable indicator hypothesis”; it posits both that exaggerated swellings signal intrinsic differences in female quality (i.e. enduring differences in phenotypic quality) and that males bias their mating behavior toward females with larger swellings (Pagel 1994). Thus, the reliable indicator hypothesis makes two main predictions: 1) males will demonstrate a preference for females with larger swellings, all else being equal; 2) superior females (with higher lifetime reproductive success) will have larger swellings. Importantly, the reliable indicator hypothesis proposes that the type of quality being signaled is a permanent characteristic of a female, and that some females are consistently superior to others, a superiority that is associated with increased lifetime reproductive success. One empirical test of this hypothesis reported support for its predictions; male baboons preferred females with larger swellings and that those females had higher infant survival (Domb & Pagel 2001). However, a reanalysis of the data presented in this study showed that it was methodologically flawed in that it failed to control for differences between baboon groups in food availability (and hence in female body condition, in the competitive environment for males, and in infant survival), which could have accounted for the observed results (Zinner et al. 2002).

Furthermore, no test of the reliable indicator hypothesis has differentiated between the permanent type of quality it proposes and the more transient type of quality that is probability of conception. Thus, without controlling for whether a given sexual cycle resulted in a conception, the reliable indicator hypothesis cannot be adequately investigated. Specifically, if conceptive swellings are larger than non-conceptive ones (which appears to be true in at least some species, (Emery & Whitten 2003; Deschner et al. 2004; Gesquiere et al. 2007; Higham et al. 2008a, 2009; Huchard et al. 2009; Fitzpatrick et al. 2014)) and, given that males of some species are able to identify and seem to prefer cycles with higher probabilities of conception (Weingrill et al. 2003; Alberts et al. 2006), then it may appear that males prefer larger swellings when, instead, they may only be tracking probability of conception. Neither Domb and Pagel (2001) nor any subsequent studies that examined the reliable indicatory hypothesis were able to control for this potential confound.

Despite the absence of resolution about what variation in swelling size can potentially signal and despite the limitations of Domb and Pagel (2001), it continues to be cited as having shown that exaggerated swellings are a reliable indicator of female quality (e.g. Gouzoules & Gouzoules 2002; Paul 2002; Jablonski 2004; Jawor et al. 2004; Dixson & Anderson 2004; Caro 2005; Preston et al. 2005; Massironi et al. 2005; Veiga & Polo 2005; Drea 2005; LeBas 2006; Weiss 2006; Polo & Veiga 2006; Pagel & Meade 2006; Gumert 2007; Watson & Platt 2008; Huchard et al. 2009; Huangshan et al. 2010; Gladbach et al. 2010; Rundle & Chenoweth 2011; Weiss et al. 2011; Haselton & Gildersleeve 2012). However, no other study has provided clear support for the reliable indicator hypothesis. One recent study found evidence for the idea that male primates prefer females with larger swellings, but was unable to control for between-cycle variation in individual females because of small sample sizes (Huchard et al. 2009).

Further, a number of studies have found that males are more responsive to conceptive cycles than to nonconceptive cycles, highlighting the importance of controlling for conceptive status in measuring male preferences (Bercovitch 1987; Bulger 1993; Weingrill et al. 2003; Setchell 2004; Alberts et al. 2006; Gesquiere et al. 2007). The one study that has successfully made this critical distinction between male response based on swelling size versus male response based on within-female changes in conception probability (between cycles) has failed to support the reliable indicator hypothesis (Deschner et al. 2004). Finally, with respect to the potential for sexual swellings to signal variation in female fitness, results are also mixed; one reports a relationship between swelling size and components of female fitness (baboons; Huchard et al. 2009) while others report an absence of one (mandrills; Setchell 2004; Setchell et al. 2006).

Thus, the functional significance of male mate choice in response to exaggerated swellings is still open. Here, we investigate the relationship between swelling size and male mating behavior as well as components of female fitness. In order to do so, we take advantage of the most detailed and largest collection to date of swelling size measurements in an ongoing study of savannah baboons (*Papio cynocephalus*; Fitzpatrick et al. 2014). This dataset has revealed critical sources of variance in swelling size; swelling size was shown to be influenced by the number of times a female had cycled since her most recent pregnancy (*cycles since resumption*), the amount of rainfall (*days since last wet month*)—which predicts food availability, and female age (Fitzpatrick et al. 2014). Together, this information accords a more precise test of the reliable indicator hypothesis than has been possible previously, by allowing us to examine male mating decisions at the same time that we control for several potential confounds. As such, we address two questions. First, do males bias their mating behavior toward females with larger swellings? Second, does variation in swelling size predict female fitness?

## METHODS

### Study Population and Data Collection

We collected morphological data (swelling size), behavioral data (male mating behavior), and proxies of female fitness from a natural population of savannah baboons (*Papio cynocephalus*) that has been under continuous study by the Amboseli Baboon Research Project (ABRP) for over four decades (Hausfater 1975; Alberts et al. 2006; Gesquiere et al. 2007; Altmann et al. 2010; Alberts & Altmann 2012). Savannah baboons exhibit female philopatry and male dispersal, live in multi-male multi-female groups where both males and females mate multiply, and breed at relatively similar frequencies year round. The Amboseli baboons occupy a short-grass savannah habitat that has undergone dramatic ecological change (from acacia woodland to more open savannah). The ecosystem is subject to extreme variation in both intra-annual and inter-annual rainfall (Altmann et al. 2002; Alberts et al. 2005).

The study population consisted of over 300 individuals of both sexes and all ages, all of which were habituated to human observers, individually identifiable by sight, and distributed across five different social groups during the period of this study. All data were collected during one field season of 12 months (November 2008 through November 2009) and a second field season of 5 months (February 2010 through June 2010).

The full analysis for this study included two discrete parts, one for each research question. Both of these analyses used morphological data as the independent variable (swelling size) and each then asked how swelling size predicted a response variable. The response variable in our first analysis was behavioral (male mate choice) and the response variables in our second analysis were life history measures (proxies of female fitness.) Because male baboons are rarely presented with an opportunity to truly choose between two simultaneously cycling females, our use of ‘male mate choice’ in this study does not necessarily imply simultaneous choice. Rather, we use two measures of male behavior to assess male mating decisions. Below, we first detail the collection of the main predictor of interest (swelling size). In subsequent sections, we describe the details of each of the two analyses, including the data collection for the response variables and the analytic approaches.

### Main Predictor Variable of Interest: Maximal Swelling Size

We used two different measures of swelling size: swelling width and swelling length. Swelling sizes were measured from digital images of individual reproductive females, collected opportunistically during daily observations. Digital images were collected using a Photoscale-2 and digital caliper, a system that allowed for the conversion of measurement in pixels to an estimate in millimeters (Fitzpatrick et al. 2014).

In order to control for the *within cycle* variation, we included only those females for whom we were able to capture maximal swelling size, designated as a size estimate that was collected within two days prior to the first day of deturgescence (“d-day”). This designation is justified by previous analyses showing that size estimates gathered on “d-2” (two days prior to deturgescence) and “d-1” (one day prior to deturgescence) were statistically equivalent (see Fitzpatrick et al. 2014 for a thorough explanation of this designation and method). In order to control for the *within individual* variation and the potential confound of conceptive status, we included only those cycles that resulted in a pregnancy. That is, this analysis was restricted only to conceptive cycles. Therefore, each female in the data set was represented only once (N = 34 females). Although spontaneous miscarriages and stillbirths occur in this population (Beehner et al. 2006; Beehner 2006), all conceptions represented in this analysis resulted in a live birth.

### Analysis 1: Male Mate Choice

#### Selection of response variables (behavioral data)

In order to assay whether male preference varied as a function of differences in swelling size between individuals, we collected behavioral data during the clear bouts of mate-guarding, or “consortships”, that take place when a female has a swelling. These are unambiguous associations between one estrous female and one adult male (Saayman 1970; Seyfarth 1978; Packer 1979; Bercovitch 1988; Alberts et al. 1996, 2003). During consortships, the consorting male attends closely to the estrous female, usually maintaining proximity and displaying vigilance. However, consortship possession is often overturned because males may fight over them intensively. As a consequence, females are not only consorted continuously during the five-day window during which they are most likely to ovulate (Wildt et al. 1977; Higham et al. 2008b; Daspre et al. 2009), but are usually consorted by more than one male during a given cycle.

We selected our behavioral response variables based on two key features of male-male competition in the presence of estrous females. First, of all the males in a social group, the highest-ranking male is the most likely to be able to exercise choice (Bulger 1993; Weingrill et al. 2003; Deschner et al. 2004; Alberts et al. 2006; Gesquiere et al. 2007). Second, consortships often attract male “following,” in which one or more males that are not the consort partner will trail the consort pair (Danish & Palombit 2014). Following males are identifiable because they clearly coordinate their movements with the consort pair, and glance at the consort pair regularly and more often than do other individuals in the group. Followers sometimes make overt attempts to take over possession of the established consortships, which usually involves charging, fighting, or coalitionary behavior with other males. Even in the absence of a clear takeover attempt, however, following behavior most likely imposes costs on the follower (e.g. limited foraging opportunities or energetic costs of vigilance, as has been documented for mate guarding itself; Alberts et al. 1996).

Thus, our two behavioral measures of male preference were 1) consortship by the highest-ranking male, and 2) proportion of consortship time that the consorting pair was trailed by at least one follower. Because males of many species bias their mating behavior toward peri-ovulatory females, we controlled for the potential effect of swelling size variation *within cycles* on male behavior by restricting our analysis to only those behavioral data that were collected during the five-day window prior to “d-day.”

#### Response variable 1: consortship by highest-ranking male

To calculate this metric, we used consortship observations that were collected on a near-daily basis as part of the ongoing ABRP data collection protocol. Consortship start time, stop time, and identity of individuals in the consorting pair, were recorded *ad libitum* whenever an observer was monitoring one of the study groups (Alberts & Altmann 2011). From these data, we created a simple binary variable for each female for whom we had a measure of maximal swelling size: the female either was or was not consorted by the highest ranking male during a given sexual cycle). We chose a binary categorization rather than a continuous measure such as a proportion (e.g., observed consort time with alpha male/total observed consort time) because when we performed a multivariate linear regression on proportion (predictors described in *Additional predictor variables* sub-subsection), the residuals were non-normally distributed, and did not respond to transformation.

#### Response variable 2: proportion of consort time with a follower

Data on followers were collected during point samples (instantaneous samples of behavior collected every two minutes) taken during 30-min focal animal samples (Altmann 1974). Focal samples were collected on consorting pairs (i.e. one estrous female and her consort partner during each focal sample). At each point sample we recorded the number of followers present and then created a proportion of consort time spent with at least one follower. Field conditions prevented the collection of focal samples from 2 of the females for which we captured maximal swelling width and length. Therefore, when using this second response variable (proportion of consort time with a follower), the sample included only 32 conceptive females. When we performed the multivariate linear regression (predictors described in *Additional predictor variables* sub-subsection), the residuals were distributed normally. Therefore, this response variable remained as a proportion in the multivariate linear regression (see Results section).

#### Additional predictor variables

In addition to swelling size, we modeled additional features of each individual female including *cycles since resumption, female age*, and *female rank*. We included one ecological variable, *days since last wet month,* and one demographic variable, the number of adult males in the queue *(queue length)*, as predictor variables (Table 1). All of these additional predictors were included to control for potentially confounding effects. The correlation coefficients between all predictor variables were less than .5 and the variance inflation factors were all less than 2.

**Table 1.** Parameters used in analysis of male mate choice as a function of swelling size.

| Predictor | Description | Range |
| --- | --- | --- |
| Swelling width | Maximal swelling width (mm) | 108.24 - 164.83 |
| Swelling length | Maximal swelling length (mm) | 121.42 – 197.74 |
| Cycles since resumption | Number of cycles since resumption | 2 - 8 |
| Female age | Female age (to nearest tenth of a year) | 7.1 - 18.3 |
| Female rank | Female dominance rank | 1 - 27 |
| Days since last wet month | Number of days since last wet month | 0 - 320 |
| Queue length | Average number of adult males in the queue<br>(number of adult males in the group - number<br>of cycling females) | 0.4 - 17 |

*Cycles since resumption* was an integer representing the number of times a female had cycled since her most recent pregnancy so that the first cycle after post-partum amenorrhea was cycle 1 and, if pregnancy did not occur, the next cycle was cycle 2, and so on. We included *cycles since resumption* because this predictor varied considerably in our data set and had a large effect on swelling size in our previous analysis (Fitzpatrick et al. 2014). That is, even though we only considered conceptive cycles in this analysis, females conceived during anywhere from 2 to 8 cycles after resumption. Therefore, we could not effectively examine the specific effect of size without controlling for an effect of *cycles since resumption*.

*Female age* was calculated to the nearest tenth of a year and was included as a predictor because age is a source of variance in swelling size in our study population (Fitzpatrick et al. 2014), indicating that we need to control for the effect of age in attempting to identify any effect of swelling size. Furthermore, fertility declines with age for female baboons (Alberts & Altmann 2003; Beehner 2006), but a study in chimpanzees found evidence that male chimpanzees prefer older females (Muller et al. 2006). That is, males may modify their investment as a function of female age, either preferring younger or older females.

*Female rank* was calculated from dyadic agonistic interactions between individuals and was represented as an ordinal number so that the highest ranking female received rank 1, the next highest ranking female was rank 2, and so on. This variable was included in our analysis because recent studies in this population found evidence of an interaction between male and female rank such that consortships were most likely when both partners were high ranking (Tung et al. 2012). In addition, some aspects of swelling size appear to be associated with female rank (Fitzpatrick et al. 2014.) Thus, we opted to control for female rank because we specifically wanted to examine the effect of swelling size on male behavior, irrespective of female rank.

*Days since last wet month* was an integer that captured the extent to which the Amboseli basin was experiencing drought or wet conditions. This predictor was especially important to account for because our two study periods spanned nearly unprecedented ecological extremes of drought (June-October 2009) and heavy rainfall (November 2009-February 2010). In addition, *days since last wet month* constrained swelling size in our previous study (Fitzpatrick et al. 2014.), and these ecological conditions may also constrain male behavior.

Finally, we included a variable representing the male competitive environment, *queue length,* as a predictor in our analysis. *Queue length* was measured as the number of adult males that were present on a given day minus the number of cycling females in the social group on that day who were within the five-day window prior to d-day. This measure of the competitive environment is based on the observation that each male can only consort one female at a time, and each female is consorted by no more than one male at a time, resulting in a competitive environment that is a function of both the number of adult males and the number of cycling females in the group. For this metric, we included cycles that were both conceptive and non-conceptive because it is extremely rare for a cycling female not to be consorted (whether or not she conceives). That is, every female that is cycling will reduce the intensity of the competitive environment. We calculated *queue length* for each of the five days during which behavioral data were collected and then took the average across that period.

#### Information theoretic analysis

Because several variables (in addition to swelling size) may influence male mating behavior, we evaluated a candidate set of models using an information theoretic approach. This method assumes that the researcher has used previous biological knowledge to select appropriate predictors, but allows for different combinations of those predictors that might be equally plausible at the outset (Burnham et al. 2011; Garamszegi 2011; Symonds and Moussalli 2011). We calculated Akaike’s Information Criterion (AIC) (Akaike 1973) for each model in our candidate set using the statistical software, R. We did this series of calculations twice: once for each response variable. We used an adjusted measure of AIC, AIC_C,_ which is recommended for smaller sample sizes (Burnham et al. 2011; Symonds & Moussalli 2011) and then calculated the Akaike weight value (*w*_i_) as a measure of goodness of fit to evaluate each of the model sets. After identifying each of the best models using the AIC approach, we used *post hoc* analyses to elucidate the effect of swelling size on our two assays of male preference.

### Analysis 2: Swellings as an Indicator of Female Fitness

To test the hypothesis that swelling size signals female fitness, we examined the relationship between variation in swelling size and proxies of female fitness. We used a classical null/alternative hypothesis testing approach for this second analysis because were wanted to explicitly test this specific hypothesis. We selected five measures, each of which is a measure of reproductive lifespan, of reproductive rate, or of early infant survival, all of which contribute ultimately to female reproductive success. Although including offspring survival as a component of parental fitness can sometimes lead to mistaken conclusions about evolutionary outcomes (Wolf & Wade 2001), we nonetheless employ the common empirical convention of using early infant survival as a reflection of female fitness. This is especially valid for long-lived primates with an extended period of dependency; in Amboseli baboons as in other wild primate populations, early infant mortality can be substantial (Altmann & Alberts 2003) and maternal characteristics contribute to infant survival (Silk et al. 2003). Therefore, males will generally benefit from mating with a female whose infants are likely to survive. Furthermore, most of our proxies have been used in previous studies (e.g., Domb and Pagel 2001). Thus, our selected measures are: 1) Age at first conception. All else being equal, individuals that mature earlier have higher lifetime fitness. In Amboseli baboons, the earliest-maturing females have on average a ½ infant advantage over their lifetime compared to the latest-maturing females (Altmann et al. 1988). 2) Survival to one year of age of the individual infant that was conceived during the cycle represented in our data set; 3) The number of a female’s infants that survived to one year of age, per reproductive year of the female’s life; and 4) The proportion of the female’s total offspring born that survived to one year of age. 5) The number of live infants born per reproductive year during the female’s life.

## RESULTS

### Analysis 1

#### Multi-model inference: predictors of whether a female was consorted by highest-ranking male on a given conceptive cycle

When we modeled the predictors of whether a female was consorted by the highest-ranking male on a given conceptive cycle, four models were produced with Δ*_i_* < 2, indicating that they had substantial and nearly equivalent support. Two of the top models included *swelling length* as a predictor variable, although none of the top models included *swelling width.* However, the effect size for *swelling length* was both weak and in the opposite direction than predicted by the reliable indicator hypothesis (ranging from −0.002 to −0.004). That is, increased swelling length decreased—rather than increased— the probability that a female was consorted by the highest-ranking male and, even then, the effect was weak. All of the top models included *number of males in the queue (queue length)*, indicating that an increased competitive environment reduced the probability that a given female would be consorted by the highest-ranking male. All but one of the top models included *cycles since resumption (csr)* as an important predictor; females who had experienced more sexual cycles since their most recent pregnancy were more likely to be consorted by the alpha male. Finally, although one of the top models included *days since last wet month (dslwm)*, the effect size was weak (0.001) (Table 2a).

**Table 2.**
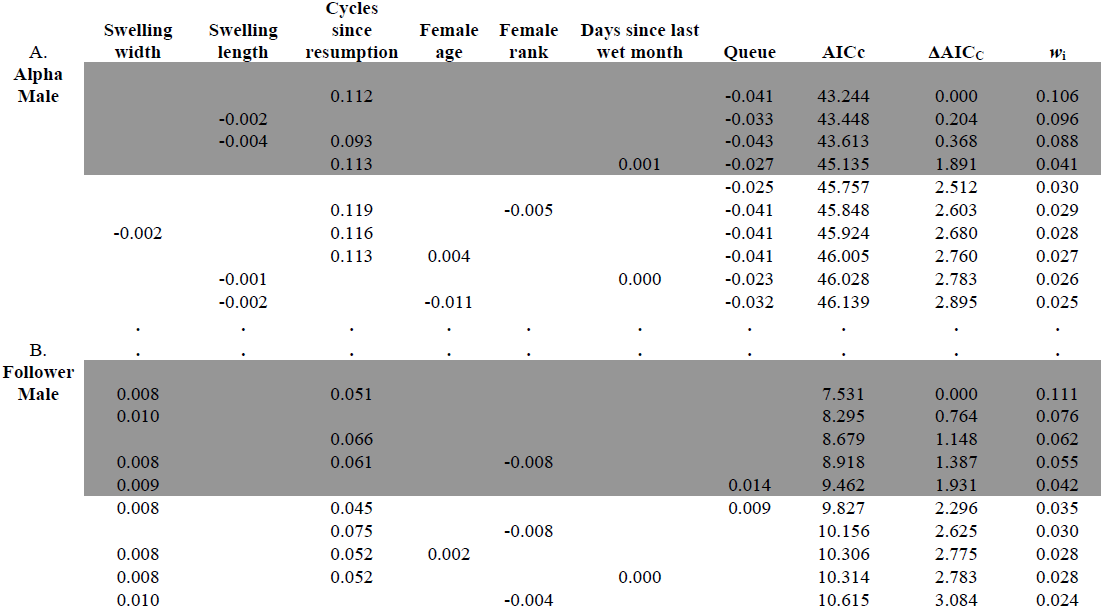
Models of fixed effects on A) the probability of being consorted by the highest-ranking male in the group (N = 34 conceptive cycles) and B) the proportion of time that a consortship was trailed by a follower male (N = 32 conceptive cycles), Shaded rows in both candidate sets indicate those models that should be considered equivalent to best model (i.e. Δ_*i*_ <2). Because only the top 10 models for each candidate set are shown, the Akaike weights that are shown (*w_i_*) do not sum to 1.

#### Multi-model inference: predictors of the proportion of time that a consorting pair had a follower

When we modeled the proportion of time that a consorting pair had a follower (considering only conceptive cycles), a total of five top models produced a Δ*_i_* < 2. In contrast to the previous analysis, *swelling width*—but not *swelling length*—predicted the proportion of time that a consortship was trailed by a follower male, and in the predicted direction (increased swelling width was associated with more time with a follower). *Swelling width* appeared in four of the five top models, and the effect size ranged from 0.008 to 0.010. *Cycles since resumption* was also an important predictor of following; it was included as a predictor in three of the five top models. Only one of the five best models included *queue length* as a predictor, indicating that the competitive environment did not influence the proportion of time that a consortship attracted a follower as much as it did the probability that a female would be consorted by the highest-ranking male. Again, one of the top models included a small effect of *days since last wet month* (Table 2b).

Taken together, the multi-model inference indicates that both measures of male preference were influenced by *cycles since resumption.* As cycle number increased, a given female was more likely to be consorted by the highest-ranking male and a given consortship was more likely to be trailed by a follower male. The effects of *queue length* and *swelling size* on male preference were more equivocal. To further examine the effects of these three variables (cycles since resumption, competitive environment, swelling size) on male mate choice, we performed *post hoc* analyses. We report only R^2^ values for these, rather than P-values, because as *post-hoc* analyses, these are subject to Type 1 errors due to multiple testing. Because the effects of *age, days since last wet month,* and *rank* were either negligible or absent, we did not investigate their effects on male mate choice any further.

#### Consortship by highest-ranking male

A bivariate analysis confirmed that females were less likely to be consorted by the highest-ranking male in groups where *queue length* was longer (i.e., where male competition was more intense; Fig. 1a). In addition, the probability of consortship by the highest-ranking male increased with the number of *cycles since resumption* (Fig. 1b) and neither measurement of swelling size predicted consortship by the alpha male in a bivariate analysis (Fig. 1c).

**Figure 1.**
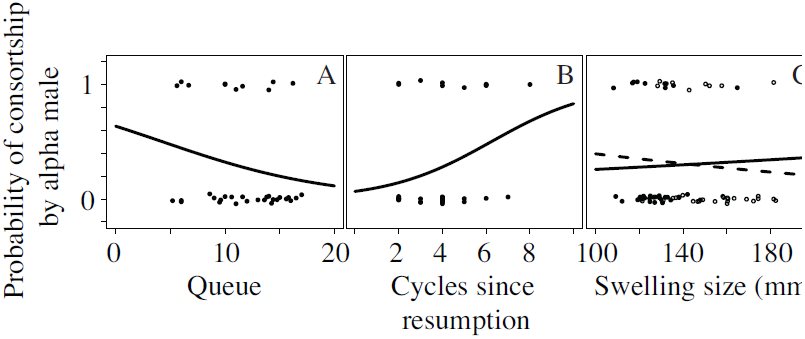
Expected probability of being consorted by the highest ranking male at any time during the five day window prior to d day a) as a function of the length of the queue, b) as a function of cycles since resumption, and c) as a function of maximal swelling size. Each trend line was generated using coefficients generated from a bivariate logistic regression with single predictor. Observed data points plotted and jittered to facilitate visibility.

#### Proportion of consort time with a follower

The proportion of time that a consortship was trailed by at least one follower was influenced both by *cycles since resumption* and *swelling width* (see above). Consequently, we performed a simple linear regression of our response variable (proportion of consort time with a follower) on each of these two predictors separately. Next, we regressed the residuals from each of these models on the other predictor, in order to isolate the effect on male behavior of each predictor independent of the other one. As both *cylces since resumption* and *swelling width* increased, consortships were more likely to attract a follower male (Fig. 2a-b; *cycles since resumption,* R^2^ = 0.06; *swelling width*, R^2^ = 0.13). That is, for a given cycle number, females with wider swellings were more likely to have a follower. Similarly, for a given *swelling width*, females who had increased *cycles since resumption* were more likely to have a follower.

**Figure 2.**
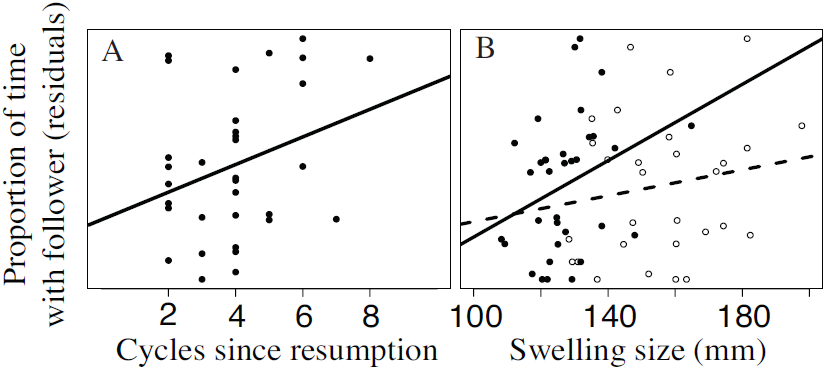
Proportion of consort time with a follower male as a function of a) cycles since resumption and b) maximal swelling size. The y axes are the residuals from a) a linear regression of *proportion of time with a follower* on *swelling width* and *swelling length* and b) a linear regression of *proportion of time with a follower* on *cycles since resumption*. 2b) Closed circles represent swelling width (solid regression line) and the open circles represent swelling length (dashed regression line).

### Analysis 2

We found no relationships between swelling size and any measure of female fitness. We fit five generalized linear models (GLM), one for each fitness proxy. In each of the five models, we included *female age* and *female dominance rank* as additional predictors to control for their potentially confounding effects on female fitness proxies. Neither maximal *swelling width* nor maximal *swelling length* were significant positive predictors of any of the five measures of female fitness (Table 3). Only the relationship between *swelling length* and *Proportion of total offspring surviving to year 1* was significant. However, in contrast to the prediction of the reliable indicator hypothesis, that coefficient (between *swelling length* and *Proportion of total offspring surviving to year 1*) was negative, indicating that, if anything, this female fitness proxy was reduced for those females with longer swellings.

**Table 3.** Coefficients and p values for relationships between swelling size (*swelling width and swelling length*) and each of the five proxies for female fitness.

| Model | Female Fitness Proxy | GLM<br>distrubtion | Coefficient |  | P Value |  |
| --- | --- | --- | --- | --- | --- | --- |
|  |  |  | Swelling<br>Width | Swelling<br>Length | Swelling<br>Width | Swelling<br>Length |
| A | Age at first conception | Poisson | 0.003 | -0.001 | 0.711 | 0.814 |
| B | Survival of infant to year 1 | Binomial | -0.068 | -0.006 | 0.225 | 0.861 |
| C | Infants born per reproductive year | Gaussian | -0.002 | 0.001 | 0.152 | 0.186 |
| D | Infants surviving to year 1 per reproductive year | Gaussian | 0.000 | -0.001 | 0.807 | 0.332 |
| E | Proportion of total offspring surviving to year 1 | Gaussian | 0.003 | **-0.004** | 0.448 | **0.045** |

## DISCUSSION

### Male Mate Choice for Cycles Since Resumption vs. Swelling Size

We examined male behavior during female sexual cycles that resulted in offspring conception, and found not only that *cycles since resumption* predicted male baboon behavior, but that it was a stronger predictor than swelling size. Specifically, females in our study who conceived after experiencing more *cycles since resumption* were both more likely to be consorted by the highest-ranking male on the conceptive cycle, and were more likely to attract at least one follower male, even when controlling for multiple potential confounds, including swelling size. In other words, cycles became more valuable from a male’s perspective as a female progressed further and further away from her previous pregnancy, perhaps because females become more physiologically able to support another pregnancy as time passes. This finding provides support for the hypothesis that males can detect the number of times a female has cycled in the recent past.

A potential adaptive explanation for this pattern emerges from the fact that each sexual cycle has some probability of *not* resulting in offspring. Thus, primate males are presented with mating opportunities that vary considerably in their fitness pay offs. Males should therefore experience selection to distinguish female sexual cycles that have a low probability of conception from those with a higher probability of conception. Indeed, previous studies both in our study population and others suggest that male baboons are able to do just that (Bercovitch 1987; Bulger 1993; Weingrill et al. 2003; Alberts et al. 2006; Gesquiere et al. 2007). Our study builds upon those results by demonstrating that males respond differently to earlier conceptive cycles (closer to a recent pregnancy) than they do to conceptive cycles that are later (further from a recent pregnancy). Indeed, our results suggest that the apparent ability of males to differentiate conceptive from non-conceptive cycles might simply reflect males’ sensitivity to the number of *cycles since resumption* that a given female has experienced, and that *cycles since resumption* may be a highly salient indicator of conception probability.

The first prediction generated by the reliable indicator hypothesis is that males should prefer females with larger swellings. That males respond to variation in *cycles since resumption* does not negate the possibility that they may also care about swelling size independently. However, our results provide only equivocal support for this first prediction. Although consortships with females that had larger swellings were more likely to attract a follower male, there was no evidence that the highest-ranking males—those most likely to exercise choice—biased their mating behavior toward females with larger swellings. An alternative interpretation of this result is that alpha males were less able to monopolize those females with larger swellings, perhaps precisely because followers challenge them more frequently. However, despite the presence of followers, high-ranking males do successfully retain consortships with females who have experienced more *cycles since resumption,* suggesting that they are, on average, able to mount the energetic resources required to maintain high-value consortships. In other words, the lack of an effect of swelling size on the behavior of alpha males suggests a lack of (or reduced) motivation rather than a lack of ability. This difference between alpha males (whose behavior was uninfluenced by swelling size) and follower males (whose behavior was) points to the possibility that alpha males and follower males may not have access to the same signal information, may prioritize that information differently, may be differently constrained, or may have different opportunities.

For instance, follower males might attend more closely than alpha males to swelling size if they tended to be newly immigrated males that would not have the information necessary to assess a female’s reproductive history or her number of *cycles since resumption* (Weingrill et al. 2003). In that case, naïve males might benefit by using swelling size as a proxy for *cycles since resumption.* However because follower males in our sample were almost entirely males who had been in the social group for longer than six months, it is unlikely that these males had less information than alpha males about the number of times that a female has cycled since her last pregnancy.

Alternatively, the differences in the costs associated with following behavior relative to those of being the primary consort partner may result in this difference in male behavior. Consorting males incur costs by forgoing foraging opportunities (Packer 1979; Alberts et al. 1996). In addition, alpha males (who are typically the most active mate guarders among adult males in a baboon group) have higher glucocorticoid levels than other males, indicating considerable energetic stress (Gesquiere et al. 2011). Therefore, successful consortship behavior requires good energetic and physiological reserves. In contrast, following behavior requires little or no special skill and the risks are presumably relatively low. Because *cycles since resumption* is likely to be a good indicator of the probability of conception, males may greatly increase their chances of capturing good reproductive opportunities (i.e. a conceptive cycle) by tracking it; furthermore, those benefits are likely to outweigh both the higher costs of being the primary consort partner and the relatively low costs of being a follower. In contrast, *swelling size* may provide some information about the quality of a reproductive opportunity (if females with larger swellings have superior physiological reserves and are therefore more likely to conceive or more able to sustain a pregnancy to term), but may not be as reliable as *cycles since resumption.* Thus, if the benefits to potential sires of larger swelling sizes are lower than the benefits of *cycles since resumption,* those benefits may not justify paying the higher costs of primary consortship but still outweigh the low costs of engaging in following behavior (see Fig 3).

**Figure 3.**
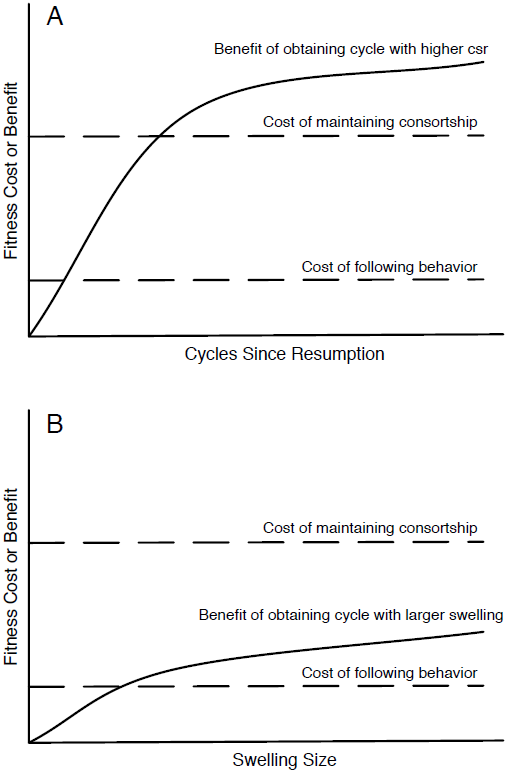
Hypothesized cost-benefit relationships that would result in the pattern observed in this study. A) Male behavior with respect to *cycles since resumption*; if the benefit to a male of monopolizing a cycle with a higher cycle number is higher than the costs of both maintaining the consortship and engaging in following behavior, then high-ranking males should work harder to obtain those consortships and follower males should invest in following them. B) Male behavior with respect to *swelling size;* if the benefit to a male of monopolizing a cycle with a larger swelling size (for a given cycle number) is higher than the cost of engaging in following behavior, but lower than the cost of maintaining the consortship, then high-ranking males should not incur the costs of obtaining those consortships, but follower males should still invest in following them.

### Competitive Environment (Number of Males in the Queue)

To our knowledge, this study is the first to control for the effect of the competitive environment (specifically the number of males relative to available females) on male mate choice with respect to swelling size. The competitive environment only affected one of our assays of male mate choice; as the number of males in the *queue* increased, the highest-ranking male was less likely to successfully monopolize a given sexual cycle. This result is consistent with previous comparative, cross-species reports (Cowlishaw & Dunbar 1991) as well as results in our study population (Alberts et al. 2003) that male ability to monopolize estrous females declines with group size. The absence of an effect of competitive environment on the behavior of follower males in the present study lends further support to the hypothesis that alpha and follower males may experience different constraints. Finally, these results highlight the fact that assays of male preference, including assays other than those we selected, may often only be meaningful if they are interpreted within the context of the competitive environment.

### Swelling Size as a “Reliable Indicator” of Fitness

Regardless of whether alpha males and follower males respond to—and respond differently to—variation in swelling size, the question remains, what information is being signaled by inter-individual variation in swelling size? Does this variation contain information that males care about, beyond probability of ovulation within a cycle and beyond the probability of conception between cycles? For the reliable indicator hypothesis to explain variation in male response to sexual swellings, variation in swelling size should predict some component of female fitness. Our analysis provides no support for the hypothesis that swelling size (either *swelling width* or *swelling length*) indicates variation in female fitness. In fact, the relationship between swelling size and female fitness was, at times, in the opposite direction opposite of that predicted by the reliable indicator hypothesis. Our study provides the strongest evidence to date that exaggerated swellings are not signals of enduring differences in fitness between females in baboons.

### Summary and future directions

We have shown that high-ranking males do not prefer females with larger swellings (when controlling for cycle number and conception) and that females with larger swellings do not have higher reproductive success. In doing so, we have tested and rejected the reliable indicator hypothesis for the function of exaggerated swellings in cercopithecine primates. Rather than tracking the potential differences in fitness between females, male baboons appear to track and target the potential for a given reproductive opportunity to result in fertilization. As the strongest such evidence to date, our results should shift the discussion about the function and evolution of exaggerated swellings from one that necessarily invokes intrinsic differences in female quality to one that focuses on temporal changes in the quality of the reproductive opportunity.

We emphasize that our test of the reliable indicator hypothesis does not address the question of whether female baboons experience sexual selection. In fact, testing the reliable indicator hypothesis cannot, in itself, identify sexual selection on female sexual swellings because the existence of male mate choice alone does not necessarily exert sexual selection pressure on female traits. Male mate choice will only exert selection pressure on swelling size if females with larger swellings receive fitness advantages as a result of being preferred beyond those that they would receive from being mated at random (see Equation 6 in Servedio 2007 for mathematical representation of this point). In other words, the common portrayal of the reliable indicator hypothesis as an explanation for the evolution of sexual swellings conflates male mate choice as a *consequence* of selection on males (if females with larger swellings have higher infant survival and therefore males prefer them) with male mate choice as a potential *cause* of selection on females. Despite this slip in logic, studies continue to cite Domb and Pagel (2001) and exaggerated swellings as an example of sexual selection in females (e.g. Paul 2002; Jawor et al. 2004; Deschner et al. 2004; Drea 2005; LeBas 2006; Fernandez & Morris 2007; Clutton-Brock 2007, 2009; Huchard et al. 2009; Rundle & Chenoweth 2011; Weiss et al. 2011; Davies et al. 2012). We caution the reader against conflating male mate choice with sexual selection on female traits. Further, in light of this caution, we highlight the importance of identifying the conditions under which male mate choice will result in selection pressure on females, versus the conditions under which male mate choice evolves but is inert as a mechanism of selection on female traits.

## ACKNOWLEDGEMENTS

We thank the Office of the President of the Republic of Kenya, the Kenya Wildlife Service and its Amboseli staff and wardens, the Amboseli-Longido pastoralist communities, Ker & Downey Safaris and Tortilis Camp for their cooperation and assistance in Kenya. This research could not have been conducted without assistance to CLF from L. Maryott and the US Embassy in Nairobi during multiple critical times. We thank the Amboseli Baboon Research Project long-term senior field researchers (R.S. Mututua, S. Sayialel, and J.K. Warutere) and the research assistants (G.Y. Marinka, C.S. Mutenkere, and B.O. Oyath) for their invaluable assistance and insight. Many people have contributed to the long-term data collection and database maintenance; in particular we thank L. Maryott, T. Fenn, and N. Learn. We thank Jeff Jacobsen for early assistance with the Photoscale-2 method, Ed Iverson and the Duke University Statistical Consulting Center for useful discussions about the statistical analysis, and Maria Servedio for helpful discussions about male mate choice. CLF was supported by Sigma Xi, Duke University Center for International Studies, Duke Biology, the Princeton Center for the Demography of Aging (P30AG024361), the Patricia William Mwangaza Foundation, the L.S.B. Leakey Foundation, an NSF Graduate Research Fellowship, and a Fulbright Fellowship. Support for the long term research project was provided by the National Science Foundation (most recently IOS 1053461 and DEB 0919200) and the National Institute of Aging (R01AG034513 and P01 AG031719). Support data from this project are available in the Dryad database: doi: XXXXX.

## LITERATURE CITED

Akaike, H. 1973. Information theory as an extension of the maximum likelihood principle. In: Second International symposium on information theory, (Ed. by B. Petrov & F. Csaki), pp. 267–281. Budapest: Akademiai Kiado.

Alberts, S. C. & Altmann, J. 2003. Matrix models for primate life history analysis. In: Primate Life History and Socioecology, (Ed. by P. Kappeler & M. E. Pereira), pp. 66–102. Chicago, Illinois: University of Chicago Press.

Alberts, S. C. & Altmann, J. 2011. Monitoring guide for the Amboseli Baboon Research Project: protocols for long-term monitoring and data collection. *Published online at* http://princeton.edu/∼baboon/monitoring_guide.html.

Alberts, S. C. & Altmann, J. 2012. The Amboseli Baboon Research Project: Themes of continuity and change. In: Long-term field studies of primates, (Ed. by P. Kappeler & D. Watts), pp. 261–288. Springer-Verlag.

Alberts, S. C. & Fitzpatrick, C. L. 2012. Paternal care and the evolution of exaggerated sexual swellings in primates. Behavioral Ecology, 23, 699–706.

Alberts, S. C., Altmann, J. & Wilson, M. L. 1996. Mate guarding constrains foraging activity of male baboons. Animal Behaviour, 51, 1269–1277.

Alberts, S. C., Watts, H. E. & Altmann, J. 2003. Queuing and queue-jumping: long-term patterns of reproductive skew in male savannah baboons, Papio cynocephalus. Animal Behaviour, 65, 821–840.

Alberts, S. C., Hollister-Smith, J. A., Mututua, R. S., Sayialel, S. N., Muruthi, P. M., Warutere, J. K. & Altmann, J. 2005. Seasonality and long-term change in a savanna environment. In: Seasonality in primates: Studies of living and extinct humans and non-human primates, (Ed. by D. K. Brockman & C. P. van Schaik), pp. 157–196. Cambridge University Press.

Alberts, S. C., Buchan, J. C. & Altmann, J. 2006. Sexual selection in wild baboons: from mating opportunities to paternity success. Animal Behaviour, 72, 1177–1196.

Altmann, J. 1974. Observational study of behaviors: Sampling methods. Behaviour, 49, 227–267.

Altmann, J. & Alberts, S. C. 2003. Variability in reproductive success viewed from a life-history perspective in baboons. American journal of human biology: the official journal of the Human Biology Council, 15, 401–9.

Altmann, J., Hausfater, G. & Altmann, S. 1988. Determinants of reproductive success in savannah baboons Papio cynocephalus. In: Reproductive Success, (Ed. by T. Clutton-Brock), pp. 403–418. Chicago: University of Chicago Press.

Altmann, J., Alberts, S. C., Altmann, S. A. & Roy, S. B. 2002. Dramatic change in local climate patterns in the Amboseli basin, Kenya. African Journal of Ecology, 40, 248–251.

Altmann, J., Gesquiere, L., Galbany, J., Onyango, P. O. & Alberts, S. C. 2010. Life history context of reproductive aging in a wild primate model. Annals of the New York Academy of Sciences, 1204, 127–38.

Andersson, M. 1994. Sexual selection. Princeton, NJ: Princeton University Press.

Beehner, J. C. 2006. The ecology of conception and pregnancy failure in wild baboons. Behavioral Ecology, 17, 741–750.

Beehner, J. C., Nguyen, N., Wango, E. O., Alberts, S. C. & Altmann, J. 2006. The endocrinology of pregnancy and fetal loss in wild baboons. Hormones and behavior, 49, 688–99.

Bercovitch, F. B. 1987. Reproductive success in male savanna baboons. Behavioral Ecology and Sociobiology, 21, 163–172.

Bercovitch, F. B. 1988. Coalitions, cooperation and reproductive tactics among adult male baboons. Animal Behavior, 36, 1198–1209.

Berglund, A., Rosenqvist, G. & Svensson, I. 1989. Reproductive success of females limited by males in two pipefish species. American Naturalist, 133, 506–516.

Bielert, C. & Anderson, C. M. 1985. Baboon sexual swellings and male response: a possible operational mammalian supernormal stimulus and response interaction. International Journal of Primatology, 6, 377–393.

Bonduriansky, R. 2001. The evolution of male mate choice in insects: a synthesis of ideas and evidence. Biological Reviews of the Cambridge Philosophical Society, 76, 305–339.

Brauch, K., Pfefferle, D., Hodges, K., Möhle, U., Fischer, J. & Heistermann, M. 2007. Female sexual behavior and sexual swelling size as potential cues for males to discern the female fertile phase in free-ranging Barbary macaques (Macaca sylvanus) of Gibraltar. Hormones and behavior, 52, 375–83.

Breaux, S. D., Watson, S. L. & Fontenot, M. B. 2012. A free choice task evaluating chimpanzees’ preference for photographic images of sex swellings: effects of color, size, and symmetry. International Journal of Comparative Psychology, 25, 118–136.

Bulger, J. B. 1993. Dominance rank and access to estrous females in male savanna baboons. Behaviour, 127, 67–103.

Burnham, K. P., Anderson, D. R. & Huyvaert, K. P. 2011. AIC model selection and multimodel inference in behavioral ecology: some background, observations, and comparisons. Behavioral Ecology and Sociobiology, 65, 23–35.

Caro, T. 2005. The adaptive significance of coloration in mammals. BioScience, 55, 125–136.

Clutton-Brock, T. 2007. Sexual selection in males and females. Science, 318, 1882–5.

Clutton-Brock, T. 2009. Sexual selection in females. Animal Behaviour, 77, 3–11.

Cowlishaw, G. & Dunbar, R. I. M. 1991. Dominance rank and mating success in male primates. Animal Behaviour, 41, 1045–1056.

Dahl, J. F., Nadler, R. D. & Collins, D. C. 1991. Monitoring the ovarian cycles of Pan troglodytes and P. paniscus: a comparative approach. American Journal of Primatology, 209, 195–209.

Danish, L. M. & Palombit, R. a. 2014. “Following,” an Alternative Mating Strategy Used by Male Olive Baboons (Papio hamadryas anubis): Quantitative Behavioral and Functional Description. International Journal of Primatology, 35, 394–410.

Daspre, A., Heistermann, M., Hodges, J. K., Lee, P. C. & Rosetta, L. 2009. Signals of female reproductive quality and fertility in colony-living baboons (Papio h. anubis) in relation to ensuring paternal investment. American journal of primatology, 71, 529–38.

Davies, N. B., Krebs, J. R. & West, S. A. 2012. An introduction to behavioural ecology.

Deschner, T., Heistermann, M., Hodges, K. & Boesch, C. 2004. Female sexual swelling size, timing of ovulation, and male behavior in wild West African chimpanzees. Hormones and behavior, 46, 204–15.

Dixson, A. F. 1983. Observations on the evolution and behavioral significance of “sexual skin” in female primates. Advances in the study of behavior, 13, 63–106.

Dixson, A. F. & Anderson, M. J. 2004. Effects of Sexual Selection Upon Sperm Morphology and Sexual Skin Morphology in Primates. International Journal of Primatology, 25, 1159–1171.

Domb, L. G. & Pagel, M. 2001. Sexual swellings advertise female quality in wild baboons. Nature, 410, 204–6.

Drea, C. M. 2005. Bateman revisited: the reproductive tactics of female primates. Integrative and comparative biology, 45, 915–23.

Edward, D. A. & Chapman, T. 2011. The evolution and significance of male mate choice. Trends in Ecology & Evolution, 26, 647–54.

Emery, M. A. & Whitten, P. L. 2003. Size of sexual swellings reflects ovarian function in chimpanzees (Pan troglodytes). Behavioral Ecology and Sociobiology, 54, 340–351.

Fernandez, A. a & Morris, M. R. 2007. Sexual selection and trichromatic color vision in primates: statistical support for the preexisting-bias hypothesis. American Naturalist, 170, 10–20.

Fitzpatrick, C., Altmann, J. & Albserts, S. 2014. Sources of variance in a female fertility signal: exaggerated estrous swellings in a natural population of baboons. Behavioral Ecology and Sociobiology,

Garamszegi, L. Z. 2011. Information-theoretic approaches to statistical analysis in behavioural ecology: an introduction. Behavioral Ecology and Sociobiology, 65, 1–11.

Gesquiere, L. R., Wango, E. O., Alberts, S. C. & Altmann, J. 2007. Mechanisms of sexual selection: sexual swellings and estrogen concentrations as fertility indicators and cues for male consort decisions in wild baboons. Hormones and behavior, 51, 114–25.

Gesquiere, L. R., Learn, N. H., Simao, M. C. M., Onyango, P. O., Alberts, S. C. & Altmann, J. 2011. Life at the top: rank and stress in wild male baboons. Science, 333, 357–60.

Gladbach, A., Gladbach, D. J., Kempenaers, B. & Quillfeldt, P. 2010. Female-specific colouration, carotenoids and reproductive investment in a dichromatic species, the upland goose Chloephaga picta leucoptera. Behavioral ecology and sociobiology, 64, 1779–1789.

Gouzoules, H. & Gouzoules, S. 2002. Primate communication: by nature honest, or by experience wise? International Journal of Primatology, 23, 821–848.

Gumert, M. D. 2007. Payment for sex in a macaque mating market. Animal Behaviour, 74, 1655–1667.

Haselton, M. G. & Gildersleeve, K. 2012. Can Men Detect Ovulation? Current Directions in Psychological Science, 20, 87–92.

Hausfater, G. 1975. Dominance and reproduction in baboons (Papio cynocephalus) - quantitative analysis. Contributions to Primatology, 7, 2–150.

Hendrickx, A. G. & Kraemer, D. C. 1969. Observations on the menstrual cycle, optiamal mating time and pre-implantaion embryos of the baboon, Papio anubis and Papio cynocephalus. Journal of Reproductive Fertility (Suppl.), 6, 119–131.

Higham, J. P., MacLarnon, A. M., Ross, C., Heistermann, M. & Semple, S. 2008a. Baboon sexual swellings: information content of size and color. Hormones and behavior, 53, 452–62.

Higham, J. P., Heistermann, M., Ross, C., Semple, S. & Maclarnon, A. 2008b. The timing of ovulation with respect to sexual swelling detumescence in wild olive baboons. Primates, 49, 295–9.

Higham, J. P., Semple, S., MacLarnon, A., Heistermann, M. & Ross, C. 2009. Female reproductive signaling, and male mating behavior, in the olive baboon. Hormones and behavior, 55, 60–7.

Huangshan, M., Zhang, M., Li, J., Zhu, Y., Wang, X. & Wang, S. 2010. Male mate choice in Tibetan macaques Macaca thibetana at. 56, 213–221.

Huchard, E., Courtiol, A., Benavides, J. A., Knapp, L. A., Raymond, M. & Cowlishaw, G. 2009. Can fertility signals lead to quality signals? Insights from the evolution of primate sexual swellings. Proceedings of the Royal Society B: Biological Sciences, 276, 1889–97.

Jablonski, N. G. 2004. The evolution of human skin and skin color. Annual Review of Anthropology, 585–623.

Jawor, J. M., Gray, N., Beall, S. M. & Breitwisch, R. 2004. Multiple ornaments correlate with aspects of condition and behaviour in female northern cardinals, Cardinalis cardinalis. Animal Behaviour, 67, 875–882.

LeBas, N. R. 2006. Female finery is not for males. Trends in ecology & evolution, 21, 283–327.

Massironi, M., Rasotto, M. B. & Mazzoldi, C. 2005. A reliable indicator of female fecundity: the case of the yellow belly in Knipowitschia panizzae (Teleostei: Gobiidae). Marine Biology, 147, 71–76.

Muller, M. N., Thompson, M. E. & Wrangham, R. W. 2006. Male chimpanzees prefer mating with old females. Current Biology, 16, 2234–8.

Nitsch, F., Stueckle, S., Stahl, D. & Zinner, D. 2011. Copulation patterns in captive hamadryas baboons: a quantitative analysis. Primates; journal of primatology, 52, 373–83.

Noe, R. & Sluijter, A. A. 1990. Reproductive tactics of male savanna baboons. Behaviour, 113, 117–170.

Nunn, C. 1999. The evolution of exaggerated sexual swellings in primates and the graded-signal hypothesis. Animal behaviour, 58, 229–246.

Olsson, M. 1993. Male preference for large females and assortative mating for body size in the sand lizard (Lacerta agilis). Behavioral Ecology and Sociobiology, 32, 337–341.

Owens, I. P. & Thompson, D. B. 1994. Sex differences, sex ratios and sex roles. Proceedings of the Royal Society B: Biological Sciences, 258, 93–9.

Packer, C. 1979. Male dominance and reproductive activity in Papio anubis. Animal behaviour, 27 Pt 1, 37–45.

Pagel, M. 1994. The evolution of conspicuous oestrous advertisement in Old World monkeys. Animal Behaviour, 47, 1333–1341.

Pagel, M. & Meade, A. 2006. Bayesian analysis of correlated evolution of discrete characters by reversible-jump Markov chain Monte Carlo. American Naturalist, 167, 808–25.

Paul, A. 2002. Sexual selection and mate choice. International Journal of Primatology, 23, 877–904.

Polo, V. & Veiga, J. P. 2006. Nest ornamentation by female spotless starlings in response to a male display: an experimental study. Journal of Animal Ecology, 75, 942–947.

Preston, B., Stevenson, I. R., Pemberton, J. M., Coltman, D. W. & Wilson, K. 2005. Male mate choice influences female promiscuity in Soay sheep. Proceedings of the Royal Society B: Biological Sciences, 272, 365–373.

Roulin, A., Jungi, T. W., Pfister, H. & Dijkstra, C. 2000. Female barn owls (Tyto alba) advertise good genes. Proceedings of the Royal Society B: Biological Sciences, 267, 937–41.

Roulin, A., Riols, C., Dijkstra, C. & Ducrest, A. 2001. Female plumage spottiness signals parasite resistance in the barn owl (Tyto alba). Behavioral Ecology, 12, 103–110.

Rundle, H. D. & Chenoweth, S. F. 2011. Stronger convex (stabilizing) selection on homologous sexual display traits in females than in males: a multipopulation comparison in Drosophila serrata. Evolution; international journal of organic evolution, 65, 893–9.

Saayman, G. S. 1970. The menstrual cycle and sexual behaviour in a troop of free ranging chacma baboons (Papio ursinus). Folia Primatologica, 12, 81–110.

Sargent, R. C., Gross, M. R. & Van Den Berghe, E. P. 1986. Male mate choice in fishes. Animal Behaviour, 34, 545–550.

Schwagmeyer, P. L. & Parker, G. A. 1990. Male mate choice as predicted by sperm competition in thirteen-lined ground squirrels. Nature, 348, 62–64.

Servedio, M. R. & Lande, R. 2006. Population genetic models of male and mutual mate choice. Evolution, 60, 674–85.

Setchell, J. M. 2004. Sexual swelling in mandrills (Mandrillus sphinx): a test of the reliable indicator hypothesis. Behavioral Ecology, 15, 438–445.

Setchell, J. M., Charpentier, M. J. E., Bedjabaga, I.-B., Reed, P., Wickings, E. J. & Knapp, L. A. 2006. Secondary sexual characters and female quality in primates. Behavioral Ecology and Sociobiology, 61, 305–315.

Seyfarth, R. M. 1978. Social relationships among male and female baboons. I. Behaviour during sexual consortship. Behaviour, 64, 204–226.

Silk, J. B., Alberts, S. C. & Altmann, J. 2003. Social bonds of female baboons enhance infant survival. Science, 302, 1231–1234.

Symonds, M. R. E. & Moussalli, A. 2011. A brief guide to model selection, multimodel inference and model averaging in behavioural ecology using Akaike’s information criterion. Behavioral Ecology and Sociobiology, 65, 13–21.

Szekely, T. & Reynolds, J. D. 1995. Evolutionary transitions in parental care in shorebirds. Proceedings of the Royal Society B: Biological Sciences, 262, 57–64.

Tung, J., Charpentier, M. J. E., Mukherjee, S., Altmann, J. & Alberts, S. C. 2012. Genetic effects on mating success and partner choice in a social mammal. The American naturalist, 180, 113–29.

Tutin, C. E. G. 1979. Mating patterns and reproductive strategies in a community of wild chimpanzees (Pan troglodytes schweinfurthii). Behavioral Ecology and Sociobiology, 6, 29–38.

Van Noordwijk, M. A. 1985. Sexual behaviour of Sumatran long-tailed macaques (Macaca fascictclaris). Zeitschrift für Tierpsychologie, 70, 277–296.

Veiga, J. P. & Polo, V. 2005. Feathers at nests are potential female signals in the spotless starling. Biology letters, 1, 334–7.

Vincent, a, Ahnesjö, I., Berglund, a & Rosenqvist, G. 1992. Pipefishes and seahorses: Are they all sex role reversed? Trends in ecology & evolution, 7, 237–41.

Watson, K. K. & Platt, M. L. 2008. Neuroethology of reward and decision-making. Philosophical transactions of the Royal Society of London. Series B, Biological sciences, 363, 3825–35.

Weingrill, T., Lycett, J. E., Barrett, L., Hill, R. A. & Henzi, S. P. 2003. Male consortship behaviour in Chacma baboons: the role of demographic factors and female conceptive probabilities. Behaviour, 140, 405–427.

Weiss, S. L. 2006. Female-specific color is a signal of quality in the striped plateau lizard (Sceloporus virgatus). Behavioral Ecology, 17, 726–732.

Weiss, S. L., Kennedy, E. A., Safran, R. J. & McGraw, K. J. 2011. Pterin-based ornamental coloration predicts yolk antioxidant levels in female striped plateau lizards (Sceloporus virgatus). The Journal of animal ecology, 80, 519–27.

Wildt, D. E., Doyle, L. L., Stone, S. C. & Harrison, R. M. 1977. Correlation of perineal swelling with serum ovarian hormone levels, vaginal cytology, and ovarian follicular development during the baboon reproductive cycle. Primates, 18, 261–270.

Wolf, J. B. & Wade, M. J. 2001. On the assignment of ^®^ tness to parents and offspring: whose ^®^ tness is it and when does it matter? 14, 347–356.

Zinner, D., Alberts, S. C., Nunn, C. & Altmann, J. 2002. Significance of primate sexual swellings. Nature, 420, 142–143.

